# Kinome state is predictive of cell viability in pancreatic cancer tumor and stroma cell lines

**DOI:** 10.1101/2021.07.21.451515

**Authors:** Matthew E. Berginski, Madison R. Jenner, Chinmaya U. Joisa, Silvia G. Herrera Loeza, Brian T. Golitz, Matthew B. Lipner, John R. Leary, Naim U. Rashid, Gary L. Johnson, Jen Jen Yeh, Shawn M. Gomez

**Affiliations:** Department of Pharmacology, The University of North Carolina at Chapel Hill, Chapel Hill, NC, USA; Lineberger Comprehensive Cancer Center, University of North Carolina at Chapel Hill, Chapel Hill, NC, USA; Department of Biostatistics, The University of North Carolina at Chapel Hill, Chapel Hill, NC, USA; Department of Surgery, University of North Carolina at Chapel Hill, Chapel Hill, NC, USA; Joint Department of Biomedical Engineering at the University of North Carolina at Chapel Hill and North Carolina State University, Chapel Hill, NC, USA; Department of Biostatistics, The University of Florida, Gainesville, FL, USA

## Abstract

Numerous aspects of cellular signaling are regulated by the kinome – the network of over 500 protein kinases that guides and modulates information transfer throughout the cell. The key role played by both individual kinases and assemblies of kinases organized into functional subnetworks leads to kinome dysregulation being a key driver of many diseases, particularly cancer. In the case of pancreatic ductal adenocarcinoma (PDAC), a variety of kinases and associated signaling pathways have been identified for their key role in the establishment of disease as well as its progression. However, the identification of additional relevant therapeutic targets has been slow and is further confounded by interactions between the tumor and the surrounding tumor microenvironment. Fundamentally, it is an open question as to the degree to which knowledge of the state of the kinome at the protein level is able to provide insight into the downstream phenotype of the cell.

In this work, we attempt to link the state of the kinome, or kinotype, with cell viability in representative PDAC tumor and stroma cell lines. Through the application of both regression and classification models to independent kinome perturbation and kinase inhibitor cell screen data, we find that the inferred kinotype of a cell has a significant and predictive relationship with cell viability. While regression models perform poorly, we find that classification approaches are able to predict drug viability effects. We further find that models are able to identify a set of kinases whose behavior in response to perturbation drive the majority of viability responses in these cell lines. Using the models to predict new compounds with cell viability effects and not in the initial data set, we conducted a validation screen that confirmed the accuracy of the models. These results suggest that characterizing the state of the protein kinome provides significant opportunity for better understanding signaling behavior and downstream cell phenotypes, as well as providing insight into the broader design of potential therapy design for PDAC.

## INTRODUCTION

While improvements in outcome for pancreatic ductal adenocarcinoma (PDAC) have occurred in the last decade with 5-year survival increasing from 5% to 10%, more options are clearly needed (Siegel et al., 2021). Major barriers to developing effective therapies have been the low tumor cellularity in PDAC and the uniquely hostile, poorly vascularized and highly fibrotic tumor microenvironment. Two tumor-intrinsic subtypes of PDAC have been identified, with the “basal-like” subtype being consistently associated with poor outcomes and potentially more resistant to the first-line cytotoxic combination FOLFIRINOX in patients (Rashid et al., 2020; Aung et al., 2018; Moffitt et al., 2015; O’Kane et al., 2020). Distinct from tumor subtypes, two molecular subtypes of PDAC stroma have been identified: “activated” which is associated with poor outcome as well as “normal” stroma. We have previously shown that the contributory cell to the “activated” stroma subtype are cancer associated fibroblasts (CAFs) and that CAFs may significantly alter response to therapy (Moffitt et al., 2015; Sherman et al., 2014; Toste et al., 2016). In addition, subtypes of CAFs have been now identified (Öhlund et al., 2014; Elyada et al., 2019). These results point to the need for new therapeutic approaches that, in addition to better activity against the tumor, also provide enhanced efficacy against the appropriate tumor microenvironment.

A number of kinases and kinase signaling pathways have been identified as components of PDAC initiation and progression, including the RAS/RAF/MAPK, p38 and PI3K pathways. More broadly, numerous aspects of cellular signaling are regulated by the kinome – the network of over 500 protein kinases that guides and modulates information transfer throughout the cell. The key role played by both individual kinases and assemblies of kinases organized into functional subnetworks leads to kinome dysregulation being a key driver of many diseases, particularly cancer. Linked with their role in disease, the druggability of kinases has led to increased interest in the development of kinase inhibitors, with over 70 now having achieved FDA approval (Cohen et al., 2021). Recent proteomics techniques including kinobeads and multiplexed inhibitor beads linked with mass spectroscopy (MIB/MS) are a relatively new mechanism which affords the ability to assess the state of the protein kinome en masse (Duncan et al., 2012; Collins et al., 2018). When combined with perturbation with targeted kinase inhibitors, the quantification of the dynamic response of the kinome provides a novel platform for the study of cell signaling and potential design of drug therapies. However, it is an open question as to what degree the behavior of the protein kinome is able to provide insight into the downstream phenotype of the cell.

As an initial attempt to assess the predictive capability of kinome behavior and potentially expand the availability of novel therapeutic options in PDAC, here we describe a modeling effort that links the measured state of the kinome, or “kinotype”, with the downstream phenotype of cell viability in PDAC relevant tumor and activated stroma cell lines. More specifically, we performed a kinase inhibitor drug screen on three patient-derived lines, with one line representing a CAF and activated stroma subtype and the other two representing the primary tumor, and measured cell viability at six doses. These PDAC-specific data were then linked with a unique public data set that assessed the broad proteomic response of the kinome to kinase inhibitor treatment via kinobeads (Klaeger et al., 2017). As a result, we were able to link cell viability in these tumor cell lines to the activation levels of 332 kinases in response to the application of 59 drugs applied at five doses.

Exploring both regression and classification approaches, we found that prediction of a drug’s effect was achievable through classification models, with ROC scores of ~0.7 for all cell lines using a random forest approach. We further use these models to identify specific kinases whose behavior in response to perturbation drove the majority of viability outcomes in these cell lines, pointing to the possibility of identifying tumor- and stroma-specific target candidates for further investigation. Finally, we experimentally validated the model by testing cell viability for previously un-tested compounds. Together, this systems view of kinome behavior and its linkage with downstream phenotypes suggests potential opportunities for improved disease classification and the identification of novel drug targets and therapeutic options in PDAC.

## METHODS

### Cell Lines

All cell lines were maintained in DMEMF12 (Gibco) supplemented with 10% FBS. All cell lines were cultured in an incubator at 37 °C with 5% CO2 and were regularly tested for mycoplasma contamination and cell line identity by short-tandem repeat testing.

### Drug screen

P0422-T1, P0411-T1, and P0119-T1 CAF cells were seeded in 384-well flat-bottom plates (Greiner) at densities of 2000, 1800, and 450 cells/well, respectively. Twenty-four hours after seeding, the 176 epigenetic and kinase inhibitor library (EpiKin176, published previously Bevill et al. (2019)) was stamped across six doses: 10 μM, 3 μM, 1 μM, 300 nM, 100 nM, 10 nM using the Biomek FXP Laboratory Automation Liquid Handling Workstation (Beckman Coulter). DMSO was used as the vehicle control at a concentration of 0.1% on cells. Synergistic effects of EpiKin176 with the combination therapy FOLFOX were assessed previously Lipner et al. (2020). Seventy-two hours post-treatment, cells were lysed with CellTiter-Glo (Promega) per the manufacturer’s protocol. Luminescence was read using the PHERAstar FS microplate reader (BMG Labtech). Data were normalized to the DMSO-only control to calculate relative viability.

### Validation studies of predicted Klaeger drugs

P0422-T1, P0411-T1, and P0119-T1 CAF cells were seeded at 3500 cells/well in white flat-bottom 96-well plates (Corning). Twenty-four hours after seeding, cells were treated with AT-13148, RGB-286638, K-252a, PF-03814735, lestaurtinib, masitinib, ripasudil, PHA-793887, AT-9283 and XL-228. Each drug was dosed at the same eight concentrations used in the Klaeger study: 30 μM, 3 μM, 1 μM, 300 nM, 100 nM, 30 nM, 10 nM and 3 nM. Seventy-two hours post-treatment, cells were lysed with CellTiter-Glo (Promega) per the manufacturer’s protocol. Luminescence was read using the PHERAstar FS microplate reader (BMG Labtech). Quality checks were performed to look at the data distribution and the presence of spatial bias on a plate. One of the replicate runs of PHA-793887 and AT-9283 failed to meet this criteria and were removed from analyses. Data were normalized to the DMSO-only (0.5% on cells) control samples on each each plate to calculate relative viability.

### General Modeling Methods

All of the models developed in this study were produced using the R programming language. The tidymodels modelling frame work was used for both the regression and binary classification models (Kuhn and Wickham, 2020). During model development we used a custom cross validation approach, which left one compound out of model training and used the remaining data for model development. For each model type tested, we also conducted a hyperparameter search over 100 combinations with latin hypercube sampling (Iman et al., 1981) across all the investigated hyperparameters. We used the the ROCR package to collect ROC and PRC values and curves (Sing et al., 2005). The functions avaible in the tidyverse package were extensively used to organize the data sets (Wickham et al., 2019).

### Cell Viability Regression Modeling

We tried three types of regression models: GLMnet (Friedman et al., 2010), random forest (Wright and Ziegler, 2017) and XGBoost (Chen and Guestrin, 2016). The hyperparameters and associated sampled ranges were as follows:

- GLMnet:

- Amount of regularization: [−10-0] with log10 transformation
- Proportion of lasso penalty: [0-1]
- Random Forest:

- Number of trees: [1000 – 5000]
- Minimum data points for split: [2 – 40]
- Number of predictors sampled: 1 or 2
- XGBoost:

- Number of trees: [1 – 2000]
- Maximum tree depth: [1-15]
- Minimum data points for split: [2 – 40]
- Reduction in loss for required for split: [−10 – 1.5] with log10 transformation
- Portion of data in fitting: [0.1 – 1]
- Number of predictors sampled: 1 or 2
- Learning rate: [−10 – −1] on log10 scale

### Cell Viability Classification Modeling

We tried three types of classification models: SVM (Karatzoglou et al., 2004), random forest (Wright and Ziegler, 2017) and XGBoost (Chen and Guestrin, 2016). We sampled over the same set of hyperparameters for the random forest and XGBoost models as in the regression models. For the SVM model, we sampled over the following hyperparameters and ranges:

- Cost of predicting prediction error: [−10-5] with log2 transformation
- Polynomial degree: [1-3]
- Kernal scaling factor: [−10 – −1] with log10 transformation

### Software and Data Availability

We have made all scripts and processing code for this project available through github under the BSD license: https://github.com/gomezlab/PDACperturbations. This repository also contains the experimental data used to build our models.

## RESULTS

### Data Organization and Distribution of Viability Values

In order to build a collection of models tailored to predicting cell viability in our PDAC cell lines, we first needed to organize and combine the data sets (Figure 1A). This was divided into two arms, the first dedicated to collecting and organizing the data from the cell screening efforts and the second to organizing the kinase activation data provided by Klaeger et al. (2017).

**Figure 1.**
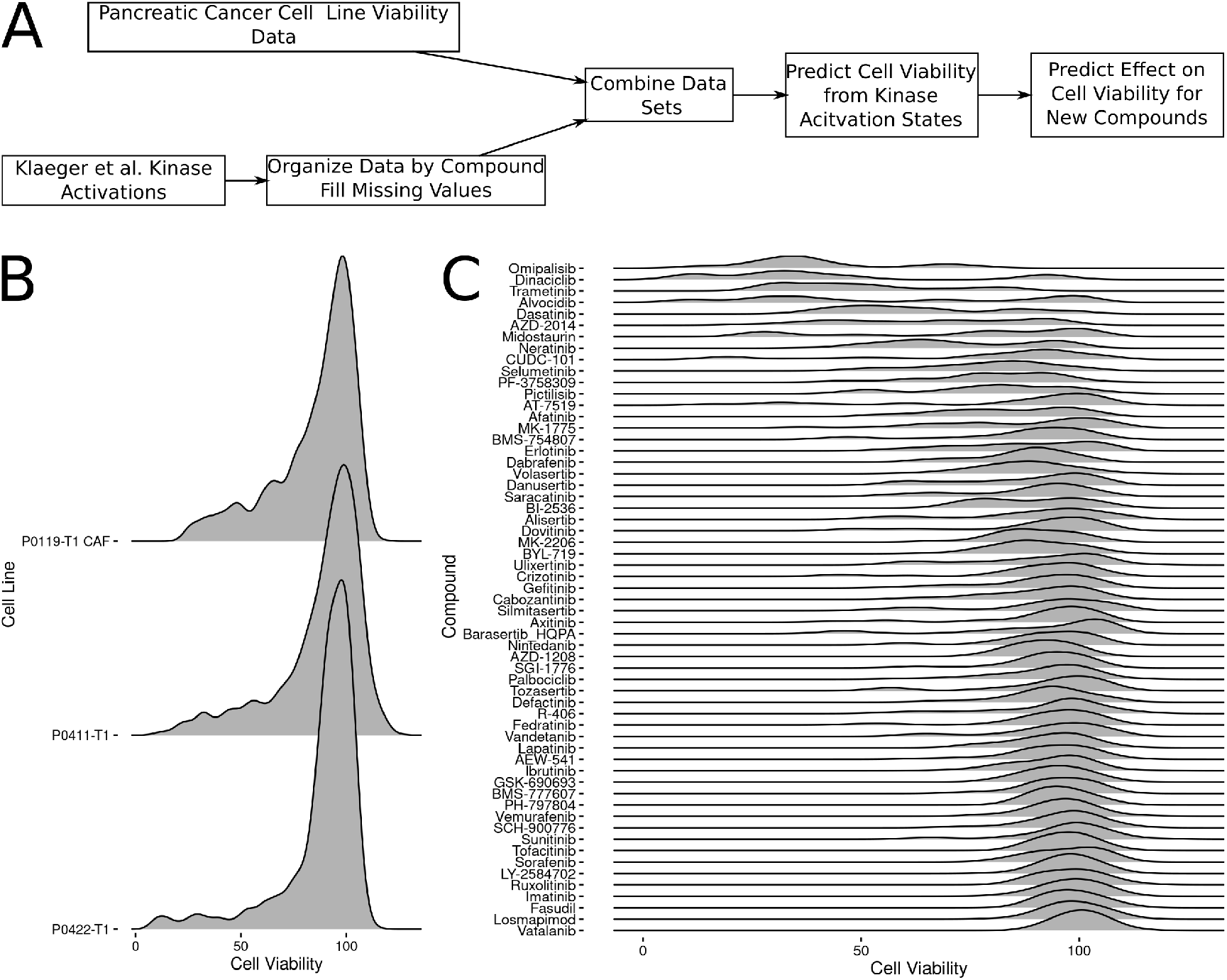
Data collection flowchart and data distribution. A: Flowchart showing data filtering, organization and downstream modeling steps. B: Distribution of cell viability values for each of the three cell lines included in this study. C: Distribution of cell viability values across the compounds overlapping between the screening data and the Klaeger et al compounds.

As the cell viability screen used a larger set of compounds than the Klaeger et al. collection during testing, the screening data was filtered to only include overlapping compounds. The Klaeger et al. collection was processed in turn, starting with the kinase activation table available in the paper supplemental materials (Table 2, Kinobeads subsheet). This spreadsheet is organized to show the kinases and a few non-kinase genes whose activation states are affected by each of the compounds tested by Klaeger et al., as such, a wide range of kinase targets are included across the range of compounds tested. For the purposes of organizing the data to enable downstream machine learning applications, we assumed that all kinases not listed with a given compound were not affected by this compound. The baseline value for an unaffected kinase is a ratio of one, so all missing compound/kinase combinations (95% of combinations) were filled in with one. We also found a small set of compound/kinase combinations where a single concentration was missing, so we filled these values in with the average value of the two closest concentrations included in the assay. Finally, we also found a few values with unexpectedly high ratio values (up to around 25), so we truncated all values in the collection to the 99.99% percentile (ratio value of 3.43).

With our two critical data sets collected and organized, we combined these data sets by matching the compounds and the corresponding concentrations. The screening data was collected at 6 compound concentrations (1e-8, 1e-7, 3e-7, 1e-6, 3e-6 and 3e-5 all in M) and all of these concentrations were matched in the Klaeger data except for 1e-5 M, which we removed from the combined data. After this filtering and matching process, we had data matches across 59 compounds. Since the full Klaeger compound set consists of 229 compounds having effects on 520 kinases, we also determined how many of these kinases were affected by our screening compound matches. This step indicated that 188 of the kinases showed zero variance in our screening compound set, so we removed these kinases from our modeling efforts. This left 332 kinases with some amount of variance in the screening compound matched data set.

The overall distribution of cell viabilities between each cell line was similar with the majority of the compounds having little effect on the cell viability (Figure 1B). This was mirrored in the distribution ofthe cell viabilities across the compounds with most of the compounds having a mean viability of above 90% (Figure 1C). Several of the compounds do demonstrate variation in the viability affects though, with alvocidib, dinaciclib and midostarin showing the greatest variability in viability across the included concentrations. The screening viability results, now matched with corresponding kinase activation states as measured in the Klaeger data, allowed us to explore the possibility of building a set of models to predict the cell viability using the kinase activation state results. We then explored the use of both regression and classification models to link changes in kinome state, as induced through kinase inhibitor treatment, with downstream cell viability as described below.

### Cell Viability Regression

With the data sets organized, we first attempted to build a set of regression models centered around predicting cell viability from the kinase activation values. We selected three model types, generalized linear models (GLMnet), random forests and XGBoost for testing. To find the optimal model for each cell line and model type, we used a cross validation approach combined with hyperparameter scanning. The cross validation approach we used was based around leaving the data pertaining to a single compound out of the training data and then building a model with the data from the remaining compounds. The primary reason we used this cross validation approach was our goal of building a model capable of predicting the cell viability from the kinase activation states for a compound which was not included in the model’s training data.

Given that each of these models have a set of hyperparameters that could effect the results of the cross validation testing, we also conducted a hyperparamter search over 100 different hyperparamter combinations for each of the three model types tested. From this collection of cross-validation predictions covering a range of possible hyperparameter sets and model types, we selected the model which minimized the root mean square error (RMSE) for each cell line and model type (Figure 2).

**Figure 2.**
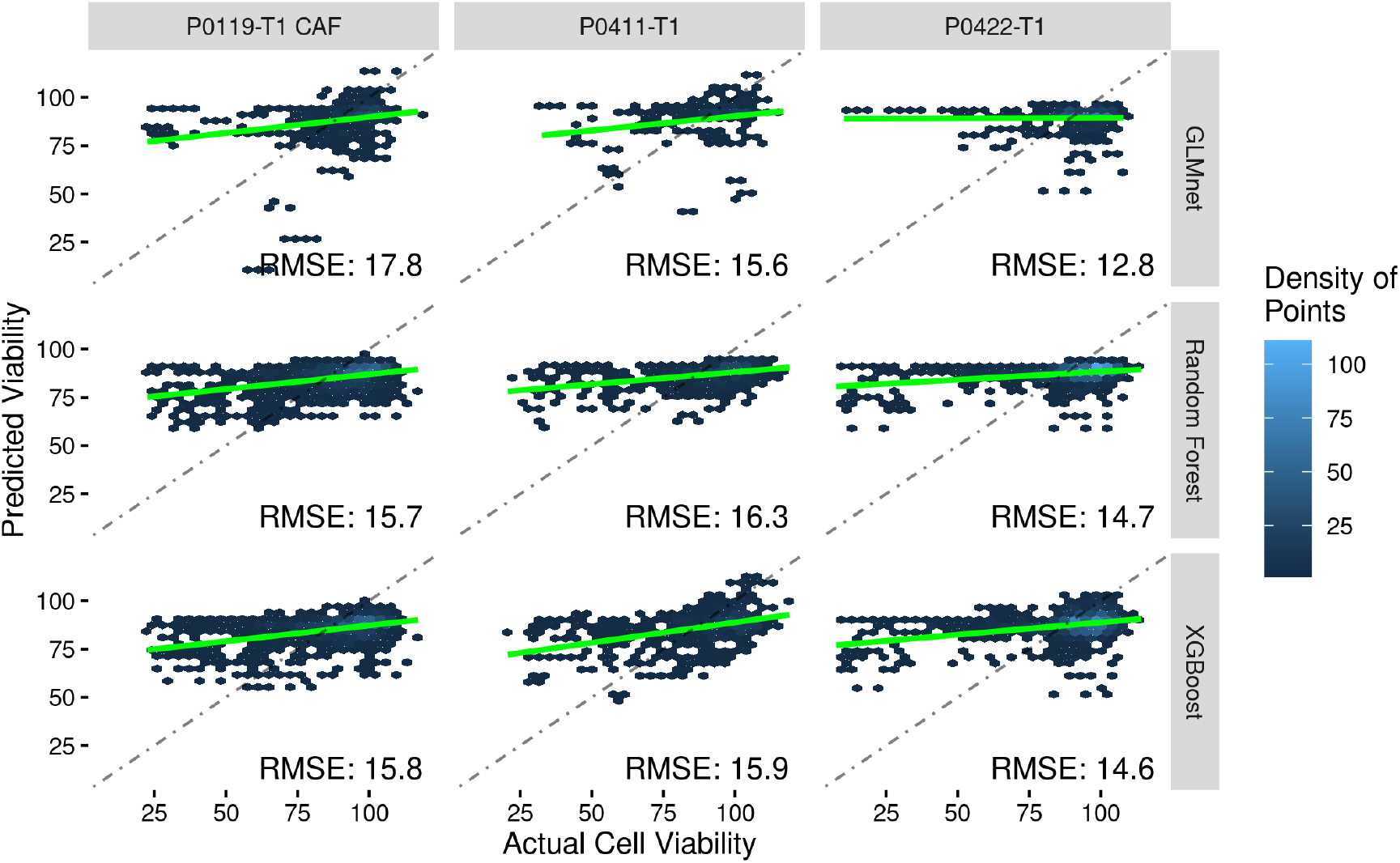
Regression models were ineffective at predicting cell viability. The results of the RMSE optimized regression models shown as hex binned heatmaps. Each column shows the optimized results for each cell line, while the rows show the type of model. The dot-dash lines show where a perfect set of predictions would appear, while the green lines show the linear best fit through the presented data points. The RMSE value is also presented in the lower corner of each plot.

Overall, none of the model types were particularly able to accurately predict the cell viability of a left out compound. This was most true for the low cell viability values in the ~25%-75% range where none of the regression methods were able to accurately predict which compound/concentration combination would result in low cell viabilities. Each of the optimized model predictions did show predictions that on average decreased for low actual cell viabilities (each green line slope is positive in Figure 2), but this reassuring result was not strong enough to reliably identify new compounds predicted to have strong effects on cell viability. While these results were disappointing, we decided to refocus our work towards a model capable of predicting a binary outcome from the cell viability results.

### Cell Viability Binary Classification

To convert the cell viability prediction problem into a binary prediction problem, we thresholded the viability values at 90% cell viability. This thresholding divided the cell the viability values into two classes with 45.5% of the CAF line data below 90% viability, 35.8% under the cutoff in the P0422-T1 line and 39% under the cutoff in the P0411-T1 line. With the data set binarized, we then experimented with building models using random forests, SVM and XGBoost. Using the same leave one compound out and hyperparameter scanning methodologies as described in the regression section, we built and tested models for each of the three cell lines (Figure 3). The best model for each modeling method, hyperparameter set and cell line was selected on the basis of the ROC score.

**Figure 3.**
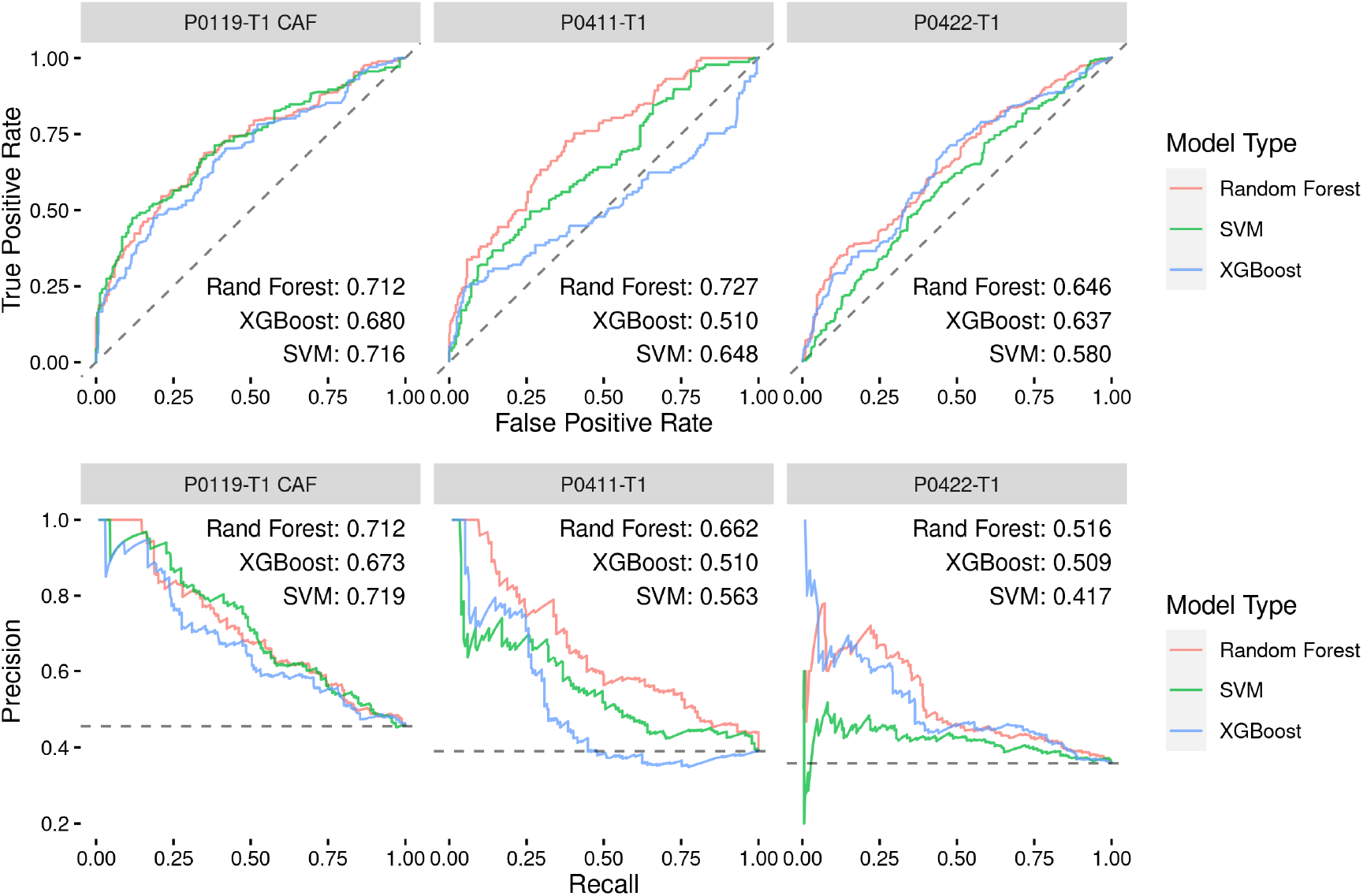
Classification models show promise in predicting drug effects on PDAC tumor and activated stroma cell lines. The top row of plots shows ROC curves for each of the best performing hyperparameter sets for each model type with the inset text indicating the area under the curve for each model type. The bottom row of plots shows the precision-recall curves for each model type with the inset text indicating the area under the curve. The cell line used to develop each model set is indicated by the gray titles above each plot.

As seen in Figure 3, classification results were significantly better than regression results, with the best single model being the random forest applied to P0411-T1 and achieving an ROC score of ~0.73. The overall ROC performance of each modelling method was similar for the P0119-T1 CAF and P0422-T1 cell lines, while the random forest method performed better in the P0411-T1 cell line. A similar pattern in performance was also observed in the precision-recall curves. This consistency of performance gave us confidence that the random forest method was most likely to yield accurate predictions when used to make predictions on compounds where we had no corresponding cell viability results. Thus, we opted to use random forest and the corresponding sets of optimal hyperparameters for each cell line to predict binarized cell viability probabilities for the compounds examined in the Klaeger set, but that were not yet tested in our cell lines.

### Using Random Forest Models to Predict Drug Effects on Cell Viability

With the random forest method and associated hyperparameters selected, we next built models for each of the cell lines using the full set of compound and concentration matched data. To examine which kinases within these models whose behavior were considered the most important for the resulting random forest models, we gathered variable importance data (Greenwell and Boehmke, 2020) for each cell line (Figure 4). These plots revealed that the top 25 kinase activation states considered important for each model was not a constant list, with only 8 kinases present in all of the top 25 lists. MAPKs (MAP2K1/2, MAP4K5) and AURKB were among the common kinase activation states across cell lines. The MAPK pathway regulates proliferation, differentiation, and apoptosis-related processes and is commonly implicated in uncontrolled tumor growth across cancers. In PDAC, the MAPKs are highly sought-after drug targets since they are downstream of KRAS-a constitutively activated protein mutated in over 90% of PDAC cases (Waters and Der, 2018). AURKs are downstream effectors in the MAPK pathway and are additionally involved with cellular processes related to mitosis, inflammation, angiogenesis, epithelial-mesenchymal transition (EMT) and metastasis. The AURKA inhibitor, alisertib, has been in clinical trials in combination with gemcitabine or nab-paclitaxel for the treatment of PDAC and other solid malignancies (NCT01924260, NCT01677559). Alisertib is expected to be investigated further in future clinical phase studies. EGFR ranked in the top 5 kinases among the PDAC lines. Overexpression of EGFR is found in 30-90% of PDAC cases and is associated with more advanced disease (Ueda et al., 2004; Tobita et al., 2003). The EGFR inhibitor, erlotinib, remains the only approved targeted therapy for PDAC. Together, these results suggest that models are identifying known, clinically-relevant kinases in PDAC that warrant further exploration as drug targets. In addition, models also identify a number of less well-studied kinases that may warrant further investigation. For instance, the serine/threonine kinase GAK (Cyclin G associated kinase) was identified within the top 6 kinases across all cell lines (Figure 4). GAK is a direct transcriptional target of p53 and has been identified as a possible serum protein biomarker for PDAC, as protein levels of GAK were found to be significantly downregulated in PDAC relative to normal pancreatic tissue in a recent multicenter trial (Gerdtsson et al., 2015). Similarly, other understudied kinases ranked in the top 25 such as CAMKK2 and PIP4K2C in P0119-T1 CAF and CSNK2A2/3 in P0422-T1 models (Oprea et al., 2018). The diversity of kinases identified as important for modeling purposes demonstrates that the role of kinases in cell viability varies even in closely related cell lines. Together, these results suggest that models are able to identify potential novel targets for further study in their role in PDAC.

**Figure 4.**
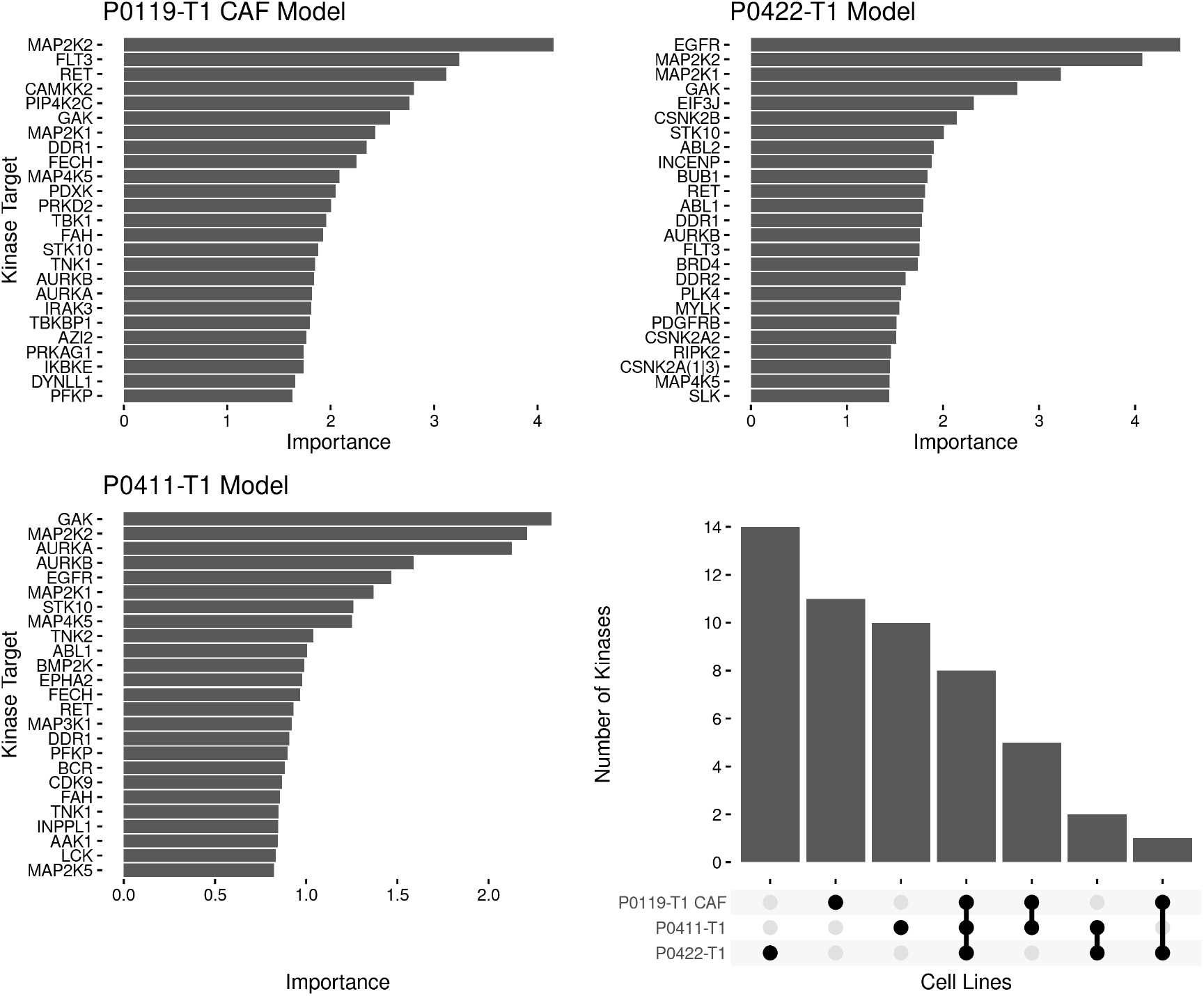
Variable importance results for optimal random forest models. A set of barplots showing the 25 kinase targets with the highest importance for each optimized random forest model and associated importance measure. The plot in the lower right corner shows an upset plot indicating the overlaps in the top 25 kinases in the three models.

With the models built for predicting the cell viability with each of our cell lines, we used these models to predict cell viability for compounds in the Klaeger et al. collection which were were previously unseen and therefore not used to build the original models. There were 165 compounds at 8 dosing concentrations in this set (1320 total combinations). Each cell-line specific model was used to predict the likelihood that a given compound and concentration combination would cause cell viability to go below 90% (Figure 5A). Overall, the P0119-T1 CAF cell line probability predictions were the highest with an average value of 0.447, followed by the P0411-T1 line at 0.390 and the P0422-T1 line at 0.371.

**Figure 5.**
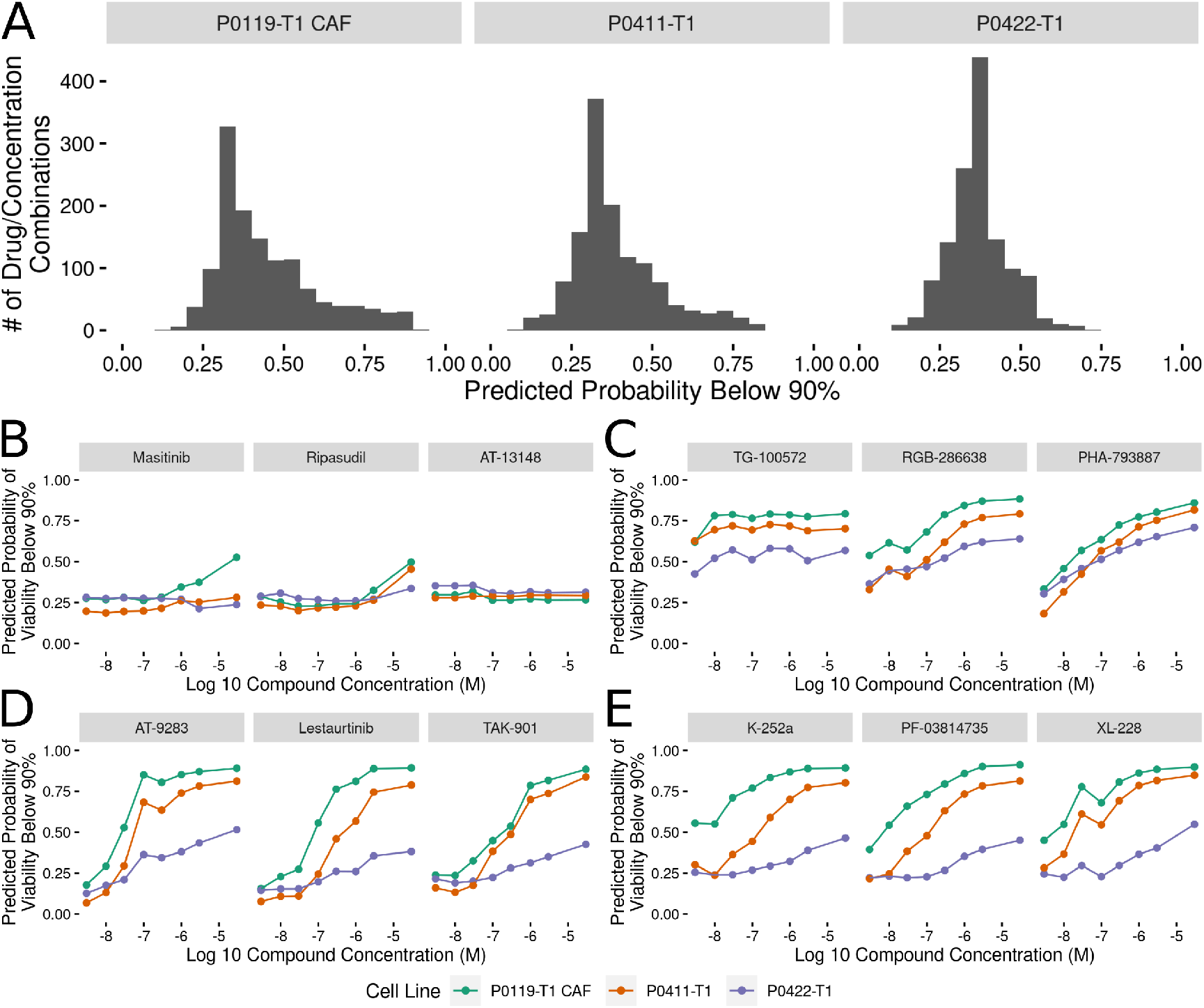
Random forest model predictions for remaining compounds. (A) Histograms showing the distributions of the probability of each compound and concentration combination having a below 90% cell viability effect. The distributions are split by cell line with the cell line name appearing in the gray title bar. (B-E) Plots showing the top 3 compound probabilities of a below 90% cell viability effect optimized for (B) the lowest average probabilities, (C) the highest average probabilities, (D) the highest range of probabilities and (E) the largest differences between the cell lines. The legend for the colors used to identify each cell line in parts B-E is presented below this collection of plots. The gray title bar in each small multiple identifies the compound.

From this collection of predictions, we extracted four categories that we thought would be especially relevant to future validation and testing efforts (Figure 5B-E). These categories of compounds are:

- Compounds with Low Predicted Effect on Viability: Predict the compounds with the lowest average probability of causing a below 90% cell viability effect (Figure 5B).
- Compounds with High Predicted Effect on Viability: Predict the compounds with the highest average probability of causing a below 90% cell viability effect (Figure 5C).
- Compounds with Large Differences across Concentrations: Predict the compounds with the highest differences in the predicted below 90% cell viability across the dose spectrum (Figure 5D).
- Compounds with Large Differences across Cell Lines: Predict the compounds with the highest average difference across the cell lines in the predicted below 90% cell viability effect (Figure 5E).

These categories of compounds optimized for a specific type of predicted effect show that the remainder of the Klaeger et al. compound set is predicted to have varying cell viability effects. Most notably, several of the compounds identified in the differences across dose curves results (Figure 5D) show large differences in the predicted effects at low compound concentration with increasing likelihood of viability effects as the compound concentration increases. This predicted effect, that mirrors a standard dose response curve, is due entirely to modifications in the kinase activation states as none of the models use compound concentration as an input value. With the model predictions collected, we selected a subset of the compounds identified in Figure 5 for follow up by cell viability testing.

### Validating a Subset of Model Predictions

To validate the prediction models, we selected 10 previously untested compounds and performed a small scale cell viability screen with the three cell lines. We conducted the validation screen at all eight of the Klaeger compound concentrations (n = 3 technical replicates, *n* = 2 biological replicates) yielding 240 cell line, compound and concentration combinations (Figure 6). We assessed these results using several different methods. First, we compared the cell viability values collected in the validation assay to the probabilities produced by each of the models and found a clear trend, with the viability values decreasing as the model probabilities increased (Figure 6A). Since the models are all predicting the probability that a given experiment will produce cell viabilities below 90%, the curve trend was in the expected downward-right direction. We also thresholded the cell viability values at 90% cell viability (as done with the training data) and produced ROC and PRC curves for the validation results (Figure 6BC). The ROC and PRC curves each demonstrate that the overall predictions for the validation compounds are performing as expected compared to the the cross validation results (Figure 6B). These global methods for quantifying model performance confirmed that the majority of the predictions made by the models were accurately observed in the followup validation screen.

**Figure 6.**
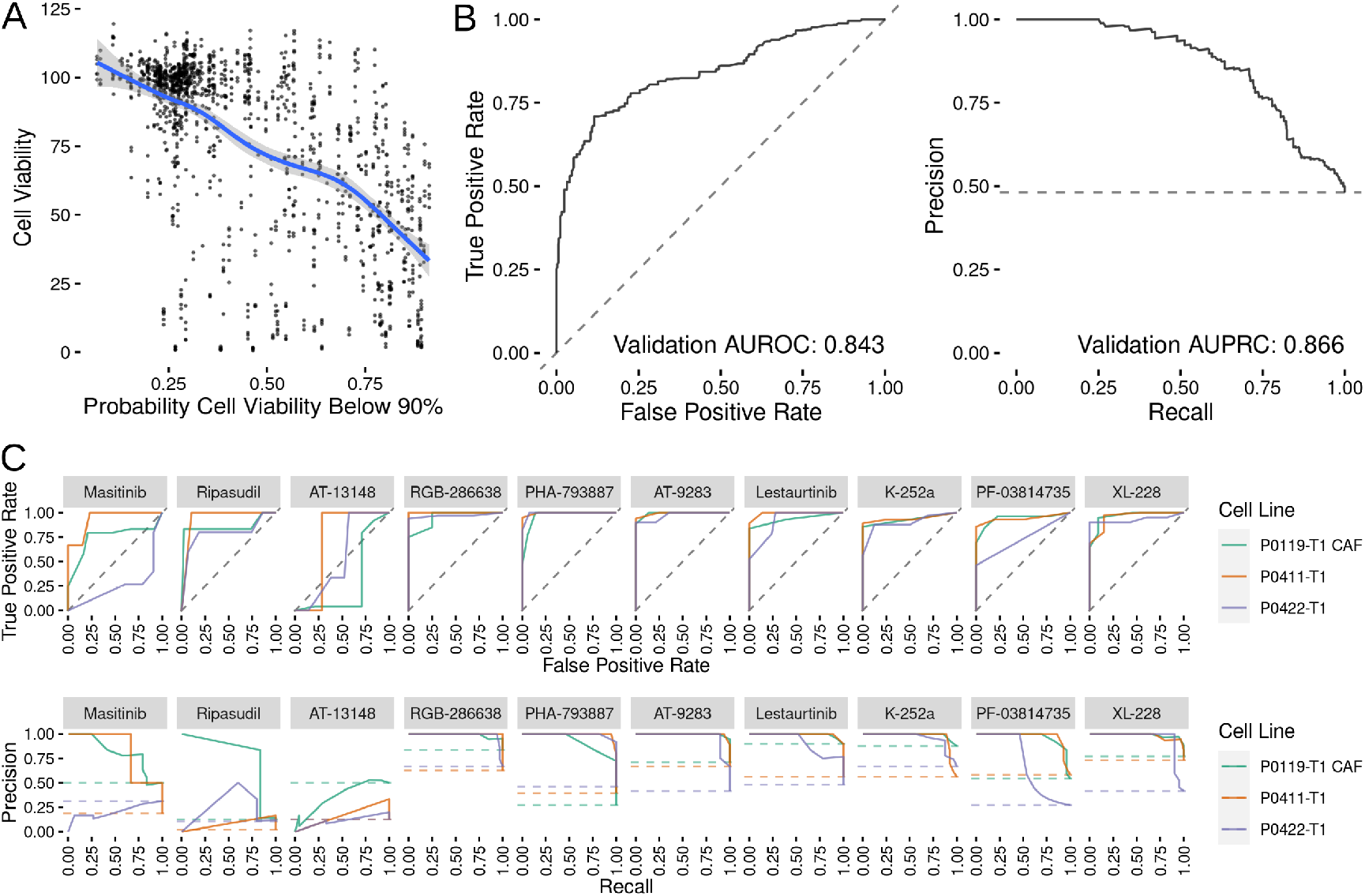
Validation results for previously untested compounds. (A) Comparison between the measured cell viability values and the predicted probability of that experiment configuration yielding a cell viability value below 90%. The blue line shows a loess fit through the data. (B) ROC and PRC curves for the validation data set thresholded at 90% viability. (C) ROC and PRC curves for all of the validation compounds tested.

We also subdivided the validation results by the tested compounds and produced individual compound ROC and PRC curve results (Figure 6C). Again, models were quite accurate for 7 out of 10 compounds tested. The models performing the worst were those predicted to have little effect on cell growth (masitinib, ripasudil and AT-13148). In the case of the compounds predicted to have a small effect, the largest effect on cell growth was observed at the highest tested compound concentration (30μM), which may be causing off target effects. Because off target effects may not be fully captured in the mass spectroscopy based assay technique used by Klaeger et al, the models would be expected to be unable to correctly predict the decrease in cell viability in these cases. With this caveat concerning interpreting the predictions at the highest concentrations in mind, we conclude that the models were largely successful at predicting the cell viability effects of a novel collection of kinase inhibitors.

## DISCUSSION

In this study, we developed a collection of models that predict cell viability from kinase activation states and used it to predict the effect of inhibitor treatment on PDAC tumor and stroma cell line models. The kinase activation data was collected through a mass spectroscopy-based method which provided an unprecedented view of how kinase inhibitors effect the entire kinome (Klaeger et al., 2017). This data has a wide range of potential uses and through this study we have connected this data to the results of a drug screening assay in PDAC cell lines. This drug screening effort overlapped with only a subset of the compounds and concentrations in the Klaeger et al. (2017) data set, but this amount of overlap was sufficient to build a collection of models which were capable of predicting the cell viability effects of kinase inhibitors. By examining the importance of each of the kinase activation states as determined by the models, we also found that a diverse set of well-studied and understudied kinase activation states were important for the modeling predictions. Of note, the dose of any compound is not used directly in the model. Rather, the generated models only use the kinase activation state data to perform inferences, with dose being indirectly encoded through the drug’s effect on kinase activation states at that dose. Using these predictions, we selected 10 additional compounds for validation screening and found that the model predictions were confirmed, with only a few exceptions. We have made all of the source code and data associated with this work publicly available through github.

While this work focuses on a small number of cell lines, the methods developed here are independent of the cell lines studied. Further work could expand the cancer types covered by gathering preliminary data using a small set of kinase inhibitors to bootstrap a set of models corresponding to any cancer type of interest. Expansion to different types of cancer would also help to clarify the role of specific kinases or collections of kinases in cell viability.

This study was also limited to assessing the role of kinase inhibitors in cell viability, but cell viability is only one of a number of possible outcome measures that could be analyzed. Any high throughput assay that can be conducted in the presence of kinase inhibitors, such as measurement of metabolite concentrations or cellular imaging assays, could be adapted to use the framework described here to attempt to generate predictive models.

Finally, since the only input to the models developed in this study is kinase activation state, it should be possible to computationally combine the activation state vectors and then make inferences about the likely cell viability results of these novel compound combinations. This would in effect be a virtual synergy screen, which could cover a much broader range of compound combinations than would be experimentally feasible. In addition, such an approach could enable the prediction of drug combinations that would preferentially effect the tumor and tumor-promoting microenvironment.

These results suggest that there is significant information encoded in the protein kinome and point to the potential to further improve predictive capabilities through the inclusion of gene expression and related data. Furthermore, this systems view of the kinome and its integration into predictive models presents opportunities for the identification of new drug targets and the design of therapies in PDAC as well as other cancers.

## ACKNOWLEDGMENTS

We would like to thank UNC research computing for access to computational resources.

